# Ground Truth–Based Evaluation of False Discovery Rate and Statistical Power in DIA Proteomics

**DOI:** 10.64898/2026.05.29.728747

**Authors:** Jay M. Yarbro, Ya Huang, Vishwajeeth Pagala, Yingxue Fu, Zhen Wang, Long Wu, Xusheng Wang, Anthony A. High, Stephanie Byrum, Junmin Peng, Zuo-Fei Yuan

## Abstract

Data-independent acquisition (DIA) mass spectrometry enables rapid proteomic quantification, yet the reliability of statistical inference in DIA-based protein quantification remains incompletely understood. Here, we systematically evaluated missingness, false discovery rate (FDR), and statistical power, defined as true positive rate (i.e. sensitivity or recall), using technical replicates and a spike-in benchmark with known ground truth. Analysis of 18 HeLa replicates revealed persistent, abundance-dependent missingness. In the spike-in experiment with five replicates, human peptides were titrated against a stable yeast background, allowing fold changes (FCs) to be compared with expected values. Across comparisons with log2FCs ranging from 0.2 to 2.5, the nominal BH-FDR substantially underestimated the true FDR. For example, at a BH-FDR threshold of 0.05, the true FDR was ∼0.2. Statistical power was ∼40% for a log2FC of 0.2 and increased to nearly 100% for a log2FC of 2.5. Additional incorporation of FC thresholds improved the true FDR for large-FC comparisons, with slight loss of power, but markedly reduced sensitivity for small-FC comparisons. Together, these results indicate that nominal FDR does not necessarily reflect actual error rates in DIA proteomics and that DIA performance is influenced by protein abundance and expected fold changes. This study provides a framework for experimental design and data interpretation in DIA-based proteomic studies.

## INTRODUCTION

Data-independent acquisition (DIA) mass spectrometry enables reproducible and high-throughput protein quantification and is increasingly used for differential proteomics studies^1-3^. Previous DIA mass spectrometry benchmarking efforts have primarily emphasized methodological and software innovations designed to enhance protein identification coverage and quantification performance^4-18^. Despite these advantages, quantitative interpretation of DIA data remains limited by missing values, variability across measurements, and uncertainty in statistical inference^19^. In particular, the relationship between nominal false discovery rate (FDR) control and actual error rates in proteomics datasets is not well studied.

Most differential proteomics analyses rely on multiple testing correction procedures, such as the Benjamini–Hochberg (BH) method^20^, to control FDR^21^. BH adjusts p-values to account for multiple comparisons and is widely used in proteomics workflows. However, it assumes that the underlying p-values are well calibrated under the null hypothesis^22, 23^. In proteomics datasets, this assumption can be difficult to satisfy. Protein-level measurements reflect multiple sources of variation, including peptide-level sampling, interference, and experimental handling, which are not explicitly modeled in standard statistical tests. As a result, nominal FDR values may not reflect the true proportion of false discoveries.

Statistical power is similarly difficult to assess in practice^24^. Power depends on effect size, variance, and sample size, but is often estimated under simplified assumptions^25, 26^. In proteomics datasets, variance is not constant^27^ and depends on protein abundance. Missing values in proteomics are also not homogeneous, arising from both stochastic sampling and abundance-dependent mechanisms^28, 29^. Low-abundance proteins are less consistently detected and show greater variability, while higher-abundance proteins are more stable. These features can affect both sensitivity and error rates, making detection performance difficult to interpret based solely on nominal statistical thresholds^30^.

To address these limitations, we generated and analyzed a technical replicate dataset and a yeast-human spike-in dataset with defined ground truth. Technical replicate analysis allows assessment of measurement variability independent of biological differences, while the spike-in design enables classification of proteins as true positives and true negatives^31^. Using this framework, we directly measured statistical power and true FDR across a range of effect sizes and significance thresholds. We show that nominal BH-FDR underestimates the true false discovery rate and that detection performance is strongly dependent on effect size. Together, these results provide an empirical basis for interpreting statistical significance in DIA proteomics and highlight limitations of relying on nominal FDR alone in differential expression analyses.

## EXPERIMENTAL SECTION

### DIA HeLa sample processing and data acquisition

200 ng of HeLa Protein Digest Standard (Pierce) were loaded on Evotips following the manufacturer’s recommendations. The samples were analyzed on a Bruker timsTOF HT instrument coupled to Evosep One (Evosep Biosystems) using a 60 samples per day (SPD) method. Evosep Performance Column (8 cm x 150 μm, C18, 1.5 μm particle size) was used for peptide separation. Data were acquired using the dia-PASEF mode^32^ with an MS1 scan range of 100–1700 m/z. Twenty-one windows were selected for serial MS2 fragmentation ranging from 475 to 1000 m/z and 0.85 to 1.27 1/K_0_, respectively.

### Human and yeast protein extraction and digestion

Human brain tissue and *S. pombe* yeast cells were lysed by bead beating and homogenized in lysis buffer containing 50 mM HEPES pH 8.5, 8 M urea, and 0.5% sodium deoxycholate. Protein concentrations were determined using the BCA assay (Thermo Fisher Scientific, 23227) and confirmed by Coomassie-stained short SDS gels^33^.

Protein samples (0.1 mg) were digested with Lys-C (1:100 w/w; Wako, 121-05063) for 2 h at 21°C. The samples were then diluted to reduce the urea concentration to 2 M, followed by overnight trypsin digestion (1:50 w/w; Promega, V5113) at 21°C. Cysteine residues were reduced by DTT and alkylated with iodoacetamide. Proteolysis was terminated by adding trifluoroacetic acid to a final concentration of 1%. The resulting peptides were desalted using Sep-Pak C18 columns (Waters) and re-quantified at UV 205 nm using a NanoDrop 2000 spectrophotometer.

### Peptide mixing and sample preparation

Human peptides were mixed with yeast peptides at defined ratios to create spike-in samples. Specifically, varying amounts of human peptides (282, 324, 400, 800, and 1600 ng) were each mixed with 2000 ng of yeast peptides in 200 µL of 0.2% formic acid to generate pooled peptide samples. Each 20 µL aliquot contained the corresponding amount of human peptides (28.2, 32.4, 40, 80, or 160 ng) and 200 ng of yeast peptides. Five technical replicates were prepared for each ratio.

### Evotip Loading

Evotips were prepared according to the manufacturer’s instructions^34^. Briefly, Evotips were wet with 20 µL of 100% acetonitrile and centrifuged at 700 × g for 60 seconds. The tips were then equilibrated with 20 µL of 0.1% formic acid in water and centrifuged (700 × g, 60 seconds). Digested peptide samples (20 µL) were loaded onto the C18 material of the Evotips by centrifugation at 700 × g for 60 seconds. For desalting, the tips were washed with 20 µL of 0.1% formic acid solution (700 × g, 60 seconds). After washing, 100 µL of 0.1% formic acid was added to each Evotip and briefly centrifuged (20 seconds) to ensure the liquid covered the C18 material and prevent drying before analysis.

### LC-MS/MS Analysis

Peptides were analyzed using an Evosep One LC system coupled to a timsTOF HT mass spectrometer^35^. The Evosep 30 SPD method was employed with an effective gradient time of 44 minutes. Peptides were separated on a 15 cm Evosep column (EV-1106, 150 µm ID, 1.9 µm C18 beads, Evosep Biosystems) at a flow rate of 500 nL/min.

Mass spectrometric analysis was performed in dia-PASEF mode^32^ with the capillary voltage set to 4 kV. The mass range was set to 100–1700 m/z for MS1 survey scans, and the ion mobility range was set to 0.60–1.60 V·s/cm^2^. The accumulation and ramp times were both set to 100 ms, resulting in a 100% duty cycle. The dia-PASEF scheme consisted of 8 PASEF frames per cycle with a total cycle time of approximately 0.9 seconds. A total of 21 isolation windows of 25 Da width were employed, covering a precursor mass range from 475 to 1000 m/z. Collision energy was ramped linearly as a function of ion mobility, from 20 eV at 1/K_0_ = 0.60 V·s/cm^2^ to 59 eV at 1/K_0_ = 1.60 V·s/cm^2^.

### DIA-NN analysis

An *in silico* spectral library was generated using DIA-NN^36^ (version 2.2.0 Academia) with the corresponding FASTAs (combined human and yeast for human-yeast mixed samples, human for HeLa samples). The parameters for the library generation were as follows: Trypsin/P with maximal 2 missed cleavages; peptide length from 7 to 30; Carbamidomethyl on C as fixed modification for human-yeast mixed samples, no fixed modification for HeLa samples; Oxidation on M as variable modification; protein N-terminal M excision on; precursor charge 1-4; precursor m/z from 400 to 1200; fragment m/z from 200 to 1800. The *in silico* spectral library was then used for the analysis of timsTOF HT data. Search and quantification parameters of DIA-NN were set as follows: precursor FDR 1%; mass accuracy at MS1 and MS2 were both set to automatic inference (0); scan window was set to automatic inference (0); MBR was turned on; Unrelated runs were turned off; protein inference was turned on and proteotypicity was set to ‘Genes’; Scoring strategy was set to ‘Peptidoforms’; machine learning was set to “NNs (cross-validated)”; quantification strategy was set to QuantUMS^37^ (high precision); cross-run normalization was turned off. The main report output by DIA-NN was processed using the following filters: Lib.Q.Value (for the respective library entry, at the precursor level) at 0.01 and Lib.PG.Q.Value at 0.01 (for the respective library entry, at the protein group level). Loading bias correction was turned on (median intensity for normalization, 15% of most variable intensities were trimmed, all proteins were normalized to yeast proteins for human-yeast mixed samples, all proteins were normalized to human proteins for HeLa samples).

### Data preprocessing and normalization

All intensity values were log_2_ transformed prior to downstream analysis^38^. Precursors shared between human and yeast were removed to retain only organism-specific signals for spike-in analyses. For protein-level quantification, precursor intensities were aggregated to protein groups as reported by DIA-NN. For the spike-in dataset, data were normalized to yeast proteins, which remain constant across conditions, to correct for loading and systematic variation. For HeLa replicate data, normalization was performed using all detected proteins. Missing values were retained during preprocessing of HeLa data to preserve the structure of incomplete detection for downstream missingness analysis. For spike-in data, missing values were imputed using MissForest^39^.

### Missingness analysis

Missing values were defined at the protein level based on incomplete detection across replicates. Protein abundance was calculated as the mean log_2_ intensity across observed values, and the probability of any missingness was modeled as a function of abundance using logistic regression in the Python package statsmodels^40^. Proteins were additionally stratified by intensity quartiles to assess abundance-dependent trends in missingness.

### Imputation benchmarking

To evaluate the impact of missing data on downstream analysis, several imputation approaches were applied to the HeLa dataset^41^. MissForest imputation was performed using a random forest–based method (100 trees, up to 10 iterations)^39^. K-nearest neighbors (KNN) imputation^42^ was performed using the R package VIM^43^ and applied with k = 5 after filtering proteins with greater than 75% missing values. Gaussian-based imputation^44^ was performed by replacing missing intensities with random draws from a normal distribution defined by a downshifted mean (μ − shift·σ) and scaled variance (width·σ), representing reconstructed low-abundance values. Samples were evaluated using relative RMSE, using the mean log_2_ intensity across observed values as a reference.

### Differential abundance analysis

Differential abundance analysis was performed at the protein level using R package limma^45^. Protein intensities were log_2_ transformed prior to analysis. Comparisons were conducted between spike-in conditions using equal replicate groups (n = 5 per condition). For each comparison, linear models were fitted to protein intensities and moderated t-statistics were computed using empirical Bayes variance estimation^45, 46^. Resulting p-values were adjusted for multiple hypothesis testing using the Benjamini–Hochberg (BH) procedure^20^ to control the false discovery rate (FDR). An initial cutoff of FDR < 0.05 was used to define protein changes with statistical significance.

### Ground truth–based statistical evaluation

Statistical performance was evaluated using the yeast–human spike-in dataset with defined ground truth. Proteins were labeled based on organism identity, with human proteins treated as true positives and yeast proteins treated as true negatives. For each comparison, proteins were classified as differentially expressed based on BH-adjusted FDR thresholds. At each threshold, true positives (TP), false positives (FP), true negatives (TN), and false negatives (FN) were determined using organism labels. Statistical power was defined as TP / (TP + FN), and true FDR was defined as FP / (TP + FP).

To characterize performance across significance thresholds, BH-adjusted FDR values were scanned across a dense range of cutoffs (∼50,000 values). For each cutoff, the corresponding confusion matrix (TP, FP, TN, FN) was computed, and power and true FDR were calculated. These values were used to evaluate the relationship between nominal FDR thresholds and observed error rates.

### Fold-change thresholding analysis

To evaluate the impact of additional effect size filtering, an absolute log_2_ fold-change threshold was applied following BH-FDR filtering. After selecting proteins with BH-FDR < 0.05, an additional cutoff was imposed based on multiples of a dataset-derived log_2_ fold-change standard deviation (k × SD). The standard deviation was defined using the log_2_ fold-change distribution of yeast proteins in the S_2_/S_1_ comparison, where no true biological change is expected, yielding a standard deviation of ∼0.25.

Thresholds corresponding to k = 0, 1, 1.5, 2, and 2.5 were evaluated. For each threshold, proteins were required to satisfy both BH-FDR < 0.05 and an absolute log_2_ fold-change greater than or equal to k × SD. Power and true FDR were then recalculated using the same ground truth framework described above, allowing assessment of the effect of fold-change filtering on detection performance across different effect sizes.

## RESULTS

### Abundance-dependent missingness limits protein detection in DIA proteomics

To define the extent and structure of missing values in DIA proteomics, we first analyzed protein identification across 18 technical replicates of the same HeLa sample (Supplementary data 1). In total, 6,286 proteins were identified across all replicates. As the number of replicates increased, cumulative protein coverage quickly approached saturation (Figure 1A). In contrast, the fraction of proteins consistently identified across all replicates progressively decreased, whereas the fraction of partially detected proteins increased. These results indicate that increasing replicate numbers improves cumulative coverage but does not eliminate incomplete protein detection.

**Figure 1.**
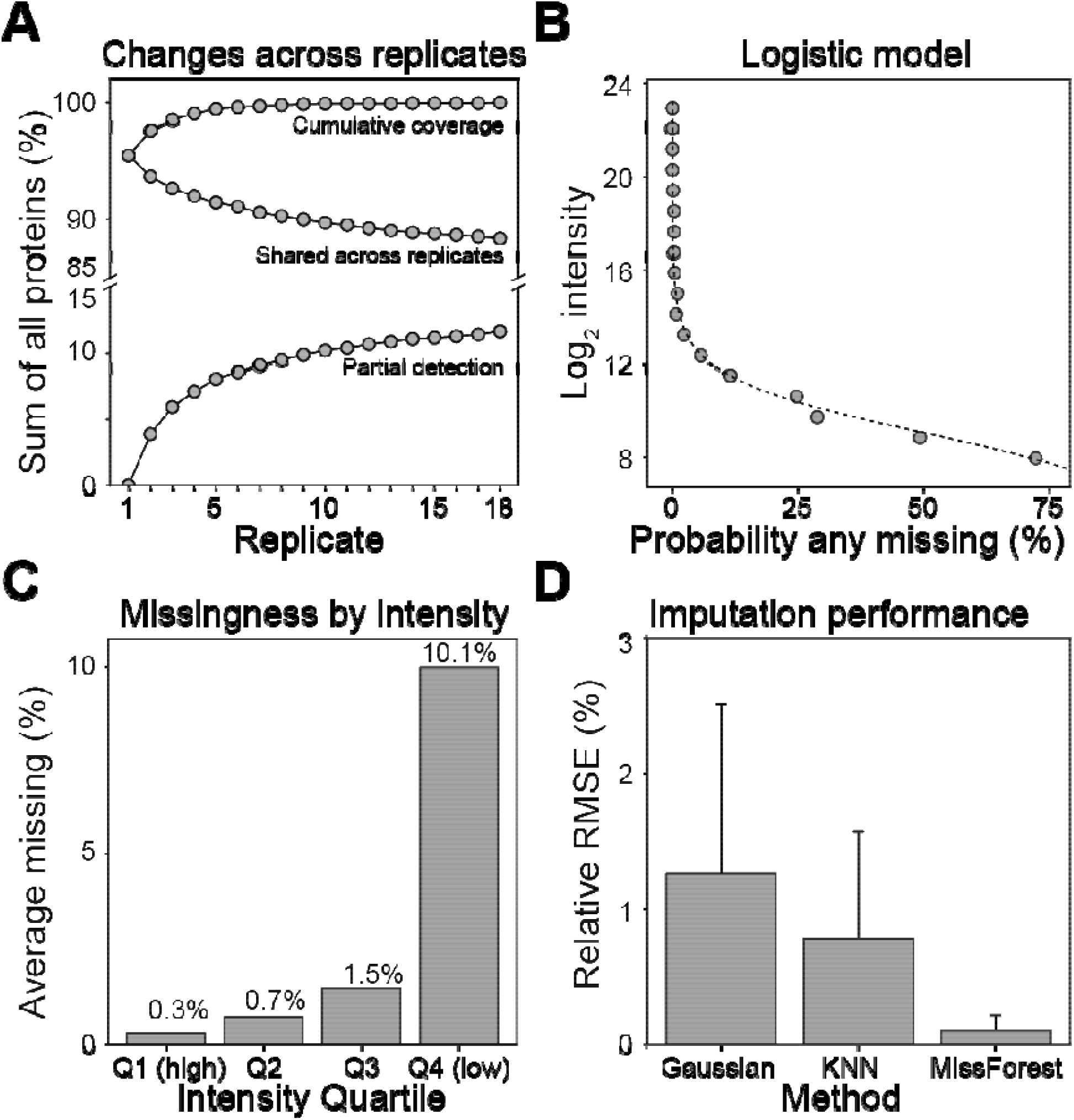
Abundance-dependent missingness and its impact on protein detection and imputation in DIA proteomics. (A) Protein identification across 18 technical replicates, showing total coverage, proteins consistently detected across all replicates, and partially detected proteins. (B) Logistic relationship between protein abundance (log_2_ intensity) and probability of any missingness. (C) Average missingness stratified by intensity quartiles. (D) Imputation performance across Gaussian, k-nearest neighbors (KNN), and MissForest methods evaluated by relative RMSE.

We next examined whether missingness was related to protein abundance. A logistic model showed a strong inverse relationship between log_2_ protein intensity and the probability of any missingness (Figure 1B), indicating that low-abundance proteins were more likely to be missed.

Consistently, stratification by intensity quartile revealed minimal missingness among the highest-abundance proteins and a marked increase in the lowest-abundance quartile (Figure 1C). Thus, missingness in DIA data is not random, but abundance dependent.

Because missing values can influence downstream quantitative analysis, we compared several common imputation strategies^39, 41^ using relative RMSE, benchmarking against the mean protein abundance from the non-imputed dataset as an estimate of the true value (Figure 1D). All methods produced relatively low relative RMSE, with MissForest (Supplementary data 2) yielding the lowest error, followed by KNN (Supplementary data 3), and Gaussian-based imputation showing the highest error among those tested (Supplementary data 4).

### Yeast–human spike-in benchmark defines ground truth and validates fold-change accuracy

To establish a ground-truth framework for evaluating statistical performance, we used a yeast– human spike-in design with a constant yeast background and increasing human peptide input across five conditions (Figure 2A). This design defines human proteins as true positives and yeast proteins as true negatives, with expected fold changes relative to the reference sample (S_1_) (Figure 2B).

**Figure 2.**
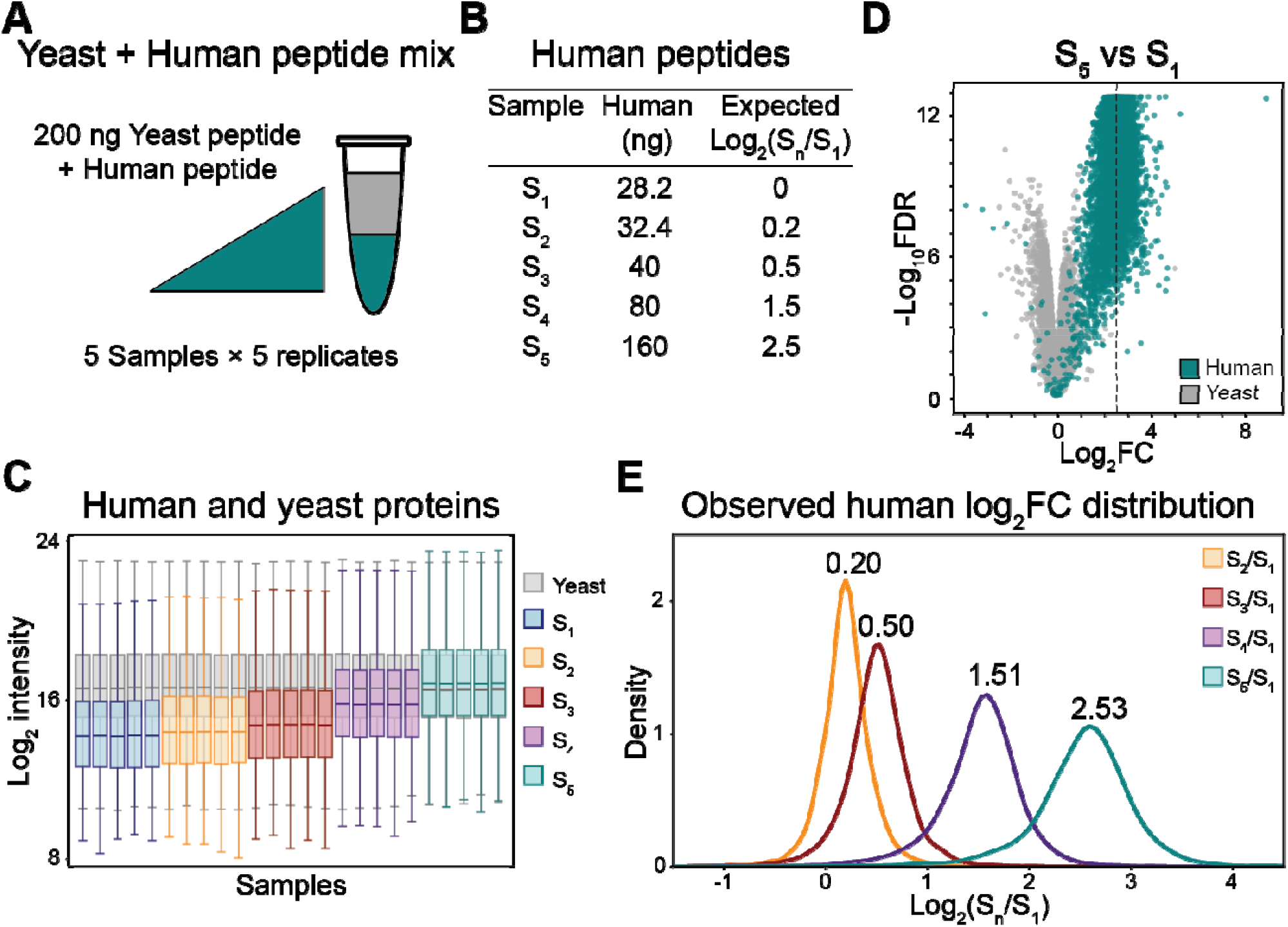
Validation of yeast–human spike-in benchmark and observed fold-change accuracy. (A) Yeast–human spike-in design with constant yeast background and increasing human input across groups (5 replicates each). (B) Human input amounts and expected log_2_(S_n_/S_1_) FCs. (C) Protein log_2_ intensity distributions across samples, with separation of stable yeast and increasing human signals. (D) Representative volcano plot of the S_5_ vs S_1_ comparison, colored by organism. Vertical dashed line corresponds to mean log_2_FC. (E) Distribution of observed human protein log_2_FCs, with mean values indicated above each distribution.

A total of 7,357 human proteins and 4,015 yeast proteins were identified. Protein intensity distributions reflected the experimental design, with yeast signals remaining stable across samples and human signals increasing with higher spike-in levels (Figure 2C, Supplementary data 5). For example, in the S_5_ versus S_1_ comparison, human proteins showed positive FCs with strong statistical significance, whereas yeast proteins were centered near zero (Figure 2D), consistent with their expected protein levels.

Human log_2_ FC distribution closely matched the expected values across all comparisons, although variability did increase with larger FCs (Figure 2E). Mean FCs aligned with the predefined inputs, indicating that measured overall fold changes were accurate. Together, these results confirm that the spike-in dataset provides a well-calibrated benchmark with known ground truth for evaluating power and false discovery rates.

### Ground truth–based evaluation of power and true false discovery rate (FDR)

Using the spike-in benchmark, we next evaluated statistical performance in terms of power and false discovery rate using known ground truth. Power was defined as the fraction of human proteins correctly identified as differentially abundant above a significance threshold, while true FDR was defined as the fraction of yeast proteins among detected proteins (Figure 3A, B).

**Figure 3.**
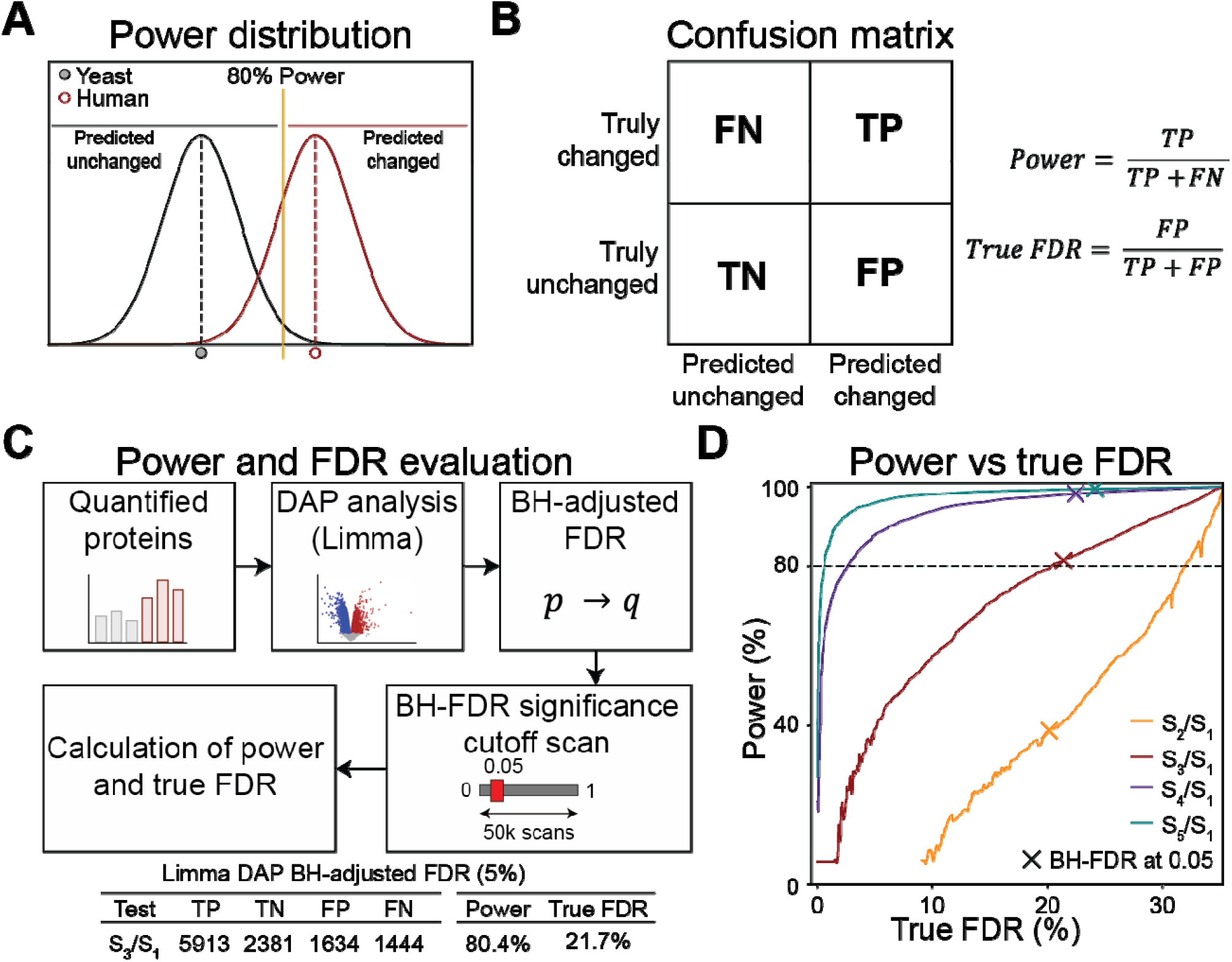
Ground truth–based evaluation of power and true false discovery rate (FDR) in DIA proteomics. (A) Conceptual illustration of statistical power relative to a significance threshold. (A) Confusion matrix defining true positives (TP), false positives (FP), true negatives (TN), and false negatives (FN), and corresponding equations for calculating power and true FDR. (C) Workflow for power and true FDR calculation following differential expression analysis, with different BH-adjusted FDR cutoffs. (D) Power versus true FDR across comparisons, with points marking the BH-FDR = 0.05 cutoff.

To quantify these metrics, we performed differential expression analysis using limma^45^ followed by Benjamini–Hochberg (BH) adjustment^20^. BH-adjusted FDR thresholds were scanned across a wide range of cutoffs, and for each cutoff, true positives, false positives, true negatives, and false negatives were computed to derive power and true FDR (Figure 3C, Supplementary data 6).

For comparisons with increasing effect sizes, power increased as expected, while true FDR decreased with more stringent thresholds (Figure 3D). However, at a commonly used threshold of BH-FDR = 0.05, true FDR were ∼0.2, substantially higher than the nominal value across all comparisons, while power varied widely depending on effect size (∼40%-99.5%). These results indicate that nominal BH-FDR does not accurately reflect true error rates in this dataset and that statistical power is strongly dependent on effect size.

### Effect of fold-change filtering on power and true false discovery rate

We next evaluated whether adding an empirical fold-change threshold improves statistical performance^47^ following BH-FDR filtering. After applying BH-FDR < 0.05, we introduced an additional absolute log_2_ fold-change cutoff defined as multiples of the standard deviation (SD) estimated from yeast S_2_/S_1_ fold-change values (where no true biological change is expected), corresponding to 0.25 (Figure 4A). Briefly, proteins were first analyzed using limma and filtered with a Benjamini–Hochberg false discovery rate (BH-FDR) cutoff of <0.05. An additional absolute log2 fold-change threshold, defined as k × SD (k = 0, 1, 1.5, 2, or 2.5), was then applied. Differentially abundant proteins were identified using these combined criteria, followed by evaluation of statistical power and true FDR.

**Figure 4.**
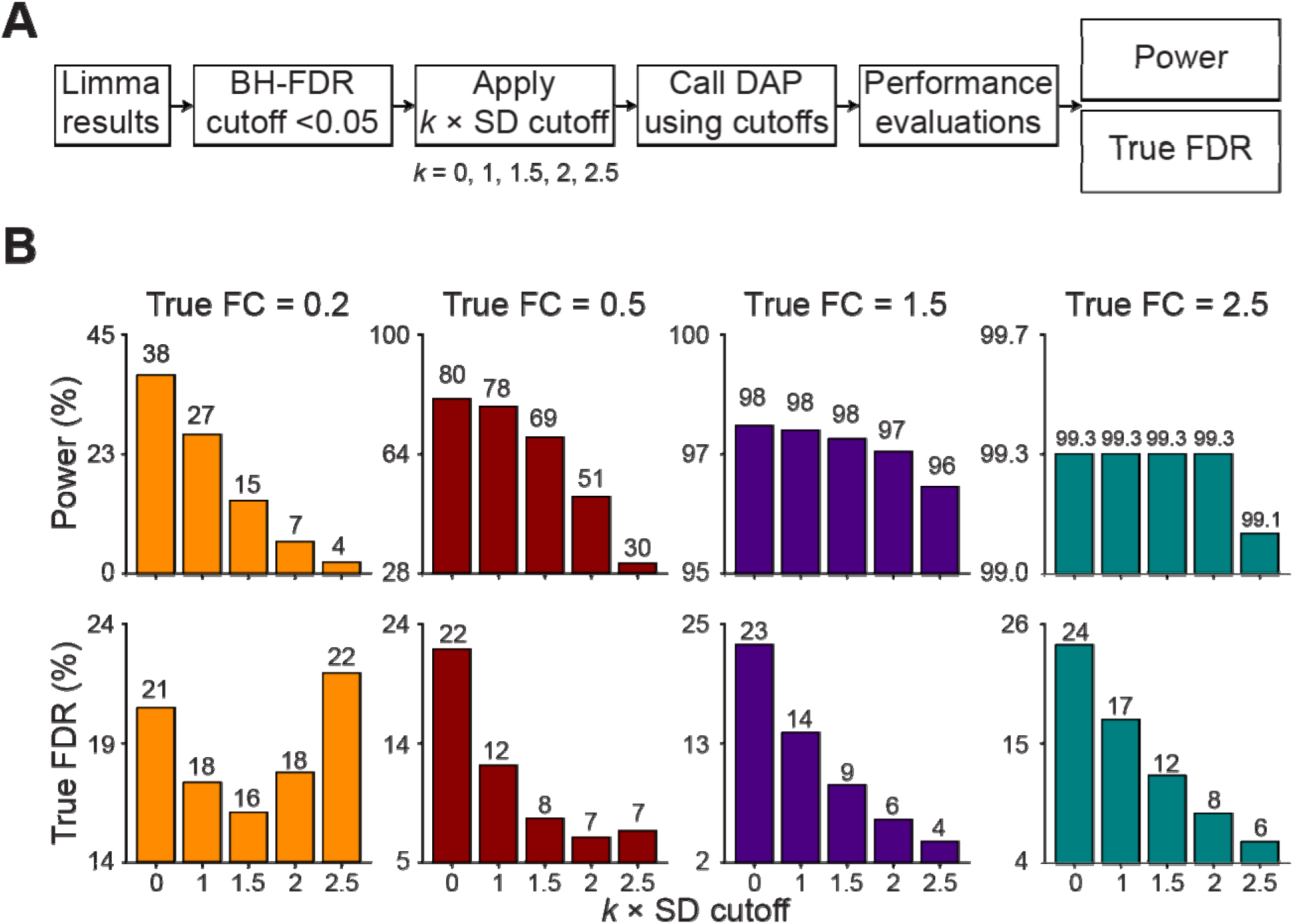
Effect of SD-based fold-change filtering on power and true false discovery rate. (A) Workflow for power and true FDR evaluation incorporating an additional absolute log_2_ fold-change cutoff (*k* × SD) following BH-FDR < 0.05. (B) Power (top) and true FDR (bottom) across increasing SD-based fold-change cutoffs (*k* = 0–2.5) for comparisons with different true effect sizes (log_2_FC = 0.2–2.5).

As shown in Figure 4B (and Supplementary Data 6), the impact of SD-based fold-change filtering depended strongly on the true effect size. For the smallest effect size (log2FC = 0.2), statistical power decreased sharply as the fold-change threshold increased, dropping from 38% at k = 0 to 27%, 15%, 7%, and 4% at k = 1, 1.5, 2, and 2.5, respectively. In contrast, the true FDR showed minimal improvement, remaining between 16% and 22%. These results indicate that additional fold-change filtering substantially reduced sensitivity for small changes without meaningfully improving error control.

For moderate effect sizes, fold-change filtering improved true FDR at the expense of reduced power. At log2FC = 0.5, power declined from 80% to 30% as k increased from 0 to 2.5, whereas true FDR improved from 22% to 7%. In contrast, filtering was more advantageous for larger effect sizes. At log2FC = 1.5, power remained high, decreasing only slightly from 98% to 97%, while true FDR decreased from 22% to 4%. Similarly, at log2FC = 2.5, power remained close to 100% across all thresholds, whereas true FDR decreased from 24% to 5%.

Together, these results demonstrate that SD-based fold-change filtering can substantially improve true FDR for large-effect comparisons with minimal loss of statistical power, but markedly reduces sensitivity when the underlying changes are small.

## DISCUSSION

Using a spike-in design with known ground truth, we show that nominal BH-FDR does not reflect the true error rate in DIA proteomics. At a threshold of 0.05, the true FDR is ∼0.2, representing an approximately four-fold underestimation. This difference is consistent across comparisons and indicates that nominal statistical control does not completely capture the error structure of the DIA proteomics data.

BH correction operates on p-values derived from quantitative measurements and assumes these p-values are well calibrated under the null hypothesis, with error arising solely from statistical variation in measured intensities^20, 48^. In practice, this assumption is incomplete. Proteomics data are generated through multiple steps, including sample handling, peptide identification, signal extraction, and protein quantification, each of which introduces sources of error that are not fully captured in the final statistical model^49, 50^. As a result, BH-FDR underestimates the true FDR because a substantial portion of the error originates outside the statistical model, leading to inaccurate p-values and a mismatch between nominal and actual error rates.

The same considerations apply to statistical power. Power is typically estimated based on replicate number, variance, and effect size^25, 51^. These estimates assume that all proteins are measured reliably and that variance is well characterized. In practice, different proteins are subject to variable detection and measurement errors. Low-abundance proteins are not consistently detected, and even when detected, their quantitative values are less stable^28^. As a result, the effective variance is higher than estimated, and power is overestimated.

This study has limitations. The analysis was performed using one workflow, and the magnitude of these effects may vary among platforms, acquisition methods, and data processing pipelines. The results were also dependent on the experimental design used here, including sample composition, replicate number, and the use of a defined biological null, which may not generalize to all proteomics studies. In addition, protein-level summarization and missing value handling can influence both variance estimates and downstream statistical testing, potentially altering the observed relationship between nominal and true error rates. Finally, proteomics data reflect a combination of biological variation and experimental error, and statistical methods applied at the final stage do not fully account for all sources of variability.

Overall, these results show that BH-FDR should not be interpreted as a direct measure of error in proteomics datasets. It reflects statistical confidence given the observed data, but underestimates errors introduced during sample handling, identification, and quantification. Introducing null experiments is a practical approach for estimating the true FDR.

## Supporting information

DIA_tables_all

## Data availability

All mass spectrometry raw data, search results, and other associated files have been deposited to the ProteomeXchange Consortium via MassIVE under the following dataset identifiers: MassIVE ID, MSV000101799; PRIDE ID, PXD078240. All data required to reproduce the analyses are provided within the manuscript and Supporting Information. Additional intermediate files and analysis scripts are available from the corresponding authors upon reasonable request.

## Supporting Information

Supplementary Data 1: Protein-level quantification matrix for 18 HeLa technical replicates (unimputed), including protein group identifiers, gene names, peptide counts, and log2-transformed intensity values (XLSX).

Supplementary Data 2: Protein-level quantification matrix following MissForest imputation, including all proteins retained for downstream analysis and corresponding log2-transformed intensity values (XLSX).

Supplementary Data 3: Protein-level quantification matrix following k-nearest neighbors (KNN) imputation (k = 5), after filtering proteins with greater than 75% missing values prior to imputation (XLSX).

Supplementary Data 4: Protein-level quantification matrix following Gaussian-based imputation using a downshifted and variance-scaled normal distribution to approximate low-abundance values (XLSX).

Supplementary Data 5: Yeast-human spike-in dataset including protein group identifiers, organism annotation, peptide counts, and log2-transformed intensity values across all spike-in conditions and replicates (XLSX).

Supplementary Data 6: Yeast-human spike-in dataset with power and true FDR calculation including comparison (test), k x SD cutoff, TP, TN, FP, FN, Power, True FDR (XLSX).

## Author Contributions

JP, JMY, and Z-FY conceived the project. YH, ZW, and VP generated mass spectrometry data. JP, AAH, XW, SH, YF, LW, Z-FY helped supervise the project. JMY, YH, and Z-FY performed the bioinformatics data analysis. JMY, YH, JP, and Z-FY wrote the manuscript with input from other authors.

## Acknowledgements

We thank all members of the Peng lab and Center for Proteomics and Metabolomics at St. Jude Children’s Research Hospital for their feedback. This work was partially supported by National Institutes of Health grants R01AG092468, RF1AG064909, U19 AG069701, and American Lebanese Syrian Associated Charities (ALSAC).

